# A high-continuity and annotated reference genome of allotetraploid Siberian wildrye (*Elymus sibiricus* L., Poaceae: Triticeae)

**DOI:** 10.1101/2024.04.17.589894

**Authors:** Jiajun Yan, Xinrui Li, Lili Wang, Daxu Li, Changmian Ji, Zujun Yang, Lili Chen, Changbing Zhang, Minghong You, Lijun Yan, Wenlong Gou, Xiong Lei, Xiaofei Ji, Yingzhu Li, Qi Wu, Decai Mao, Dan Chang, Shangang Jia, Ping Li, Jianbo Zhang, Yanli Xiong, Yi Xiong, Mengli Han, Zhao Chen, Xinchao Cheng, Juan Tang, Wengang Xie, Wenhui Liu, Hongkun Zheng, Xiao Ma, Xuebing Yan, Shiqie Bai

## Abstract

*Elymus sibiricus* L. (Siberian wildrye, *Es*), a species belonging to the wheat tribe, is extensively employed as forage and for the reclamation of degraded grasslands within the Qinghai-Tibet Plateau (QTP). This study provides a high-quality reference genome assembly for the allotetraploid *Es*, which is composed of 14 pseudomolecules with the total genome size of 6.57 Gb. Our finding suggest that large-scale bursts of retrotransposons are critical for the genome expansion of *Es*. We discovered a translocation event between the *Es*4H and *Es*6H chromosomes with a low frequency of combination. Phylogenetic analyses of 90 *Es* accessions and 25 diploid accessions representing proposed ancestors from various habitats revealed the existence of four distinct populations. We further provided support for the hypothesis that the QTP is the center of origin and genetic diversity for *Es*. Collectively, our study offers valuable insights into the evolution of *Es*, as well as providing genomic resources for genetic enhancement in the *Elymus* genus and wheat tribe.

## Introduction

Heteropolyploidization is a crucial mechanism facilitating plant adaptation to varying environmental conditions ^[1]^. *Elymus* L. (*Elymus* sensu lato), the largest and most widely spread genus in the wheat tribe (Triticeae, Poaceae), consists of around 150 perennial allopolyploid species. Each *Elymus* s.l. species contains the St genome sourced from the *Pseudoroegneria* genus, along with genomes from diverse Triticeae genera like *Hordeum* (H genome), *Agropyron* (P genome), *Australopyrum* (W genome), and an unidentified donor (Y genome) ^[2,3]^. The genomic composition of *Elymus* s.l. species makes them particularly valuable for agricultural breeding and for investigating the mechanisms of polyploidization and tolerance to environmental stress. Previous studies, predominantly relying on organelle genes/markers or single-copy nuclear genes, have shown that the St subgenome across *Elymus* s.l. species has a polyphyletic origin ^[4]^, and comparable results were also observed in diverse geographical populations of *Elymus sibiricus* when analyzing data from complete chloroplast genome ^[5]^. However, the current research is limited to specific genomic segments, which hinders a comprehensive coverage and corroborates the reliability of this conclusion. As of now, genomic resources for *Elymus* s.l. remain still limited, with only one St whole-genome sequence currently available ^[6]^. This limitation highlights the critical need for extensive genomic datasets to accurately interpret the evolutionary history and resilience mechanisms of *Elymus* s.l.

The allotetraploid *E. sibiricus* (2n = 4X = 28, StStHH), the model species of the *Elymus* s.l., commonly referred to as Siberian wildrye, is a perennial, self-pollinating grass native to the alpine regions of the Qinghai-Tibet Plateau (QTP) and northern Asia ^[7,8]^. It naturally thrives in a variety of habitats, including alpine meadows, forest glades, scrub, mountain slopes, and screes in river valleys, at altitudes ranging from 1500 to 4900 m ^[9]^. Leveraging its adaptability to harsh alpine environments, high productivity, excellent nutritional quality, and prolonged growing season, *E. sibiricus* has been extensively utilized for revegetation of degraded grassland, soil stabilization and forage production in the QTP and other high-altitude areas ^[10]^. The remarkable phenotypic diversity and substantial genetic variation observed across various wild eco-geographical populations or accessions indicate that *E. sibiricus* has likely developed and preserved genetic adaptations through natural selection to thrive in diverse environmental conditions and habitats ^[11]^. Therefore, it serves as an ideal candidate species for exploring stress resistance genes and investigating the mechanisms underlying the species’s adaptation to extreme conditions. Recent research has shifted focus towards revealing genetic variations related to adaptation by employing novel methodologies such as genome-environment association (GEA) analysis and selective sweep analysis. Nevertheless, the lack of comprehensive genome-wide data poses a significant obstacle to the study of *E. sibiricus* and its related species, constraining the investigation of their genetic foundations for adapting to varying environments.

In this study, we employed next-generation sequencing, single-molecular real-time (SMRT) sequencing, and chromatin conformation capture (Hi-C) technologies to generate a high-quality chromosome-level genome of the cultivar *E. sibiricus* ‘Chuancao No. 2’. Our findings not only provide insights into the evolution history and environmental adaptation of *E. sibiricus*, but also may contribute to the development of genomics-assisted breeding strategies for enhancing stress tolerance in this crucial grass and other Triticeae species.

## Results and Discussion

### High-quality genome assembly

A reference genome assembly was generated for the allotetraploid *E. sibiricus* cv. ‘Chuancao No. 2’ (hereafter referred to as *Es*), an elite Chinese grass cultivar recognized for its high yield and excellent resistance. Flow cytometry analysis indicated that the estimated genome size of *Es* is 6.74 Gb, aligning with the previously reported genome size of 6.86 Gb from k-mer frequency analyses ^[12]^ (Table 1, Supplementary Fig. 1). By combining the 639.99 Gb (97.34×) Pacbio long reads (Supplementary Table 1) and 469.18 Gb (68 X) Illumina short reads, a 6.57 Gb assembly with a contig N50 of 4.46 Mb was generated, covering 97.48% of the estimated genome size (Table 1, Supplementary Table 1 and 2). This assembly was further improved by utilizing 461.17 Gb of Hi-C data, resulting in the anchoring of 99.28% of the assembly (6.53 Gb) into 14 pseudomolecules, achieving a scaffold N50 value of 460.93 Mb (Fig. 1A and Supplementary Fig. 2; Table 1, and Supplementary Table 3a and 3b).The pseudomolecules were subsequently divided into the St (3.15 Gb) and H (3.37 Gb) subgenomes (hereafter referred to as *Es*St and *Es*H), based on their sequence identity to the diploid possible progenitor H genome from barley (Supplementary Table 3c). The strong collinear relationships between the physical positions of the pseudomolecules and the previously reported genetic linkage map ^[13]^ suggests the accurate phasing of the *Es*St and *Es*H subgenomes (Supplementary Fig. 3). Remapping of Illumina shorted reads revealed an average alignment rate of 99.82% to the assembly (Supplementary Table 4). In addition, 93.33% of the 242,078 unigenes identified from the transcriptome data were accurately aligned to a single contig within the genome assembly using BLAT ^[14]^ (Supplementary Table 5). The long terminal repeat (LTR) assembly index (LAI) of this assembly is 13.71, meeting the reference quality criterion ^[15]^ (Supplementary Table 6). Benchmarking Universal Single-Copy Orthologs (BUSCO) analyses showed 99.13% (1600 of 1,614) assembly completeness ^[16]^ (Supplementary Table 7). The potential centromeric regions of 14 chromosomes were identified in *Es* using centromeric elements from Cereba and Quinta in the wheat tribe ^[17]^ (Supplementary Fig. 4 and Supplementary Table 8). These results suggest a highly accurate and near-complete assembly of *E. sibiricus*.

**Table 1.** Statistics of *Es* genome assembly and annotation.

| Genomic feature |  | Value | Percentage (%) |
| --- | --- | --- | --- |
| Genomic assembly | Estimated genome size | 6.74 Gb |  |
|  | Assembled genome size | 6.57 Gb |  |
|  | Assembled genome percentage | 97.48% |  |
|  | Sequence anchored to chromosomes | 6.53 Gb | 99.28 |
|  | Number of scaffolds | 5,692 |  |
|  | Longest scaffold | 492,218,085 bp |  |
|  | N50 scaffold length | 460,926,496 bp |  |
|  | Total size of assembled contigs | 6,574,523,438 bp |  |
|  | Number of contigs | 8,060 |  |
|  | Longest contig | 28,011,202 bp |  |
|  | N50 contig length | 3,808,844 bp |  |
|  | GC content | 45.68% |  |
| TE | Number of Retrotransposons | 3,410,831 | 72.65 |
|  | Length of Retrotransposons | 4,776,281,916 bp |  |
|  | Number of DNA transposons | 1,194,947 | 12.99 |
|  | Length of DNA transposons | 854,283,427 bp |  |
|  | Number of unclassified TE | 169,370 | 1.26 |
|  | Length of unclassified TE | 83,001,403 bp |  |
|  | Number of total repeat sequences | 4,775,148 | 86.90 |
|  | Length of total repeat sequences | 5,713,566,746 bp |  |
| Genome annotation | Number of gene models (High confidence) | 79,002 |  |
|  | Number of gene models (Lower confidence) | 22,767 |  |
|  | Number of noncoding RNAs | 8,820 |  |
|  | Gene length mean | 6,384.51 bp |  |
|  | CDS length mean | 1,315.85 bp |  |
|  | Exon length mean | 314.6 bp |  |
|  | Intron length mean | 1,271.24 bp |  |

**Fig 1.**
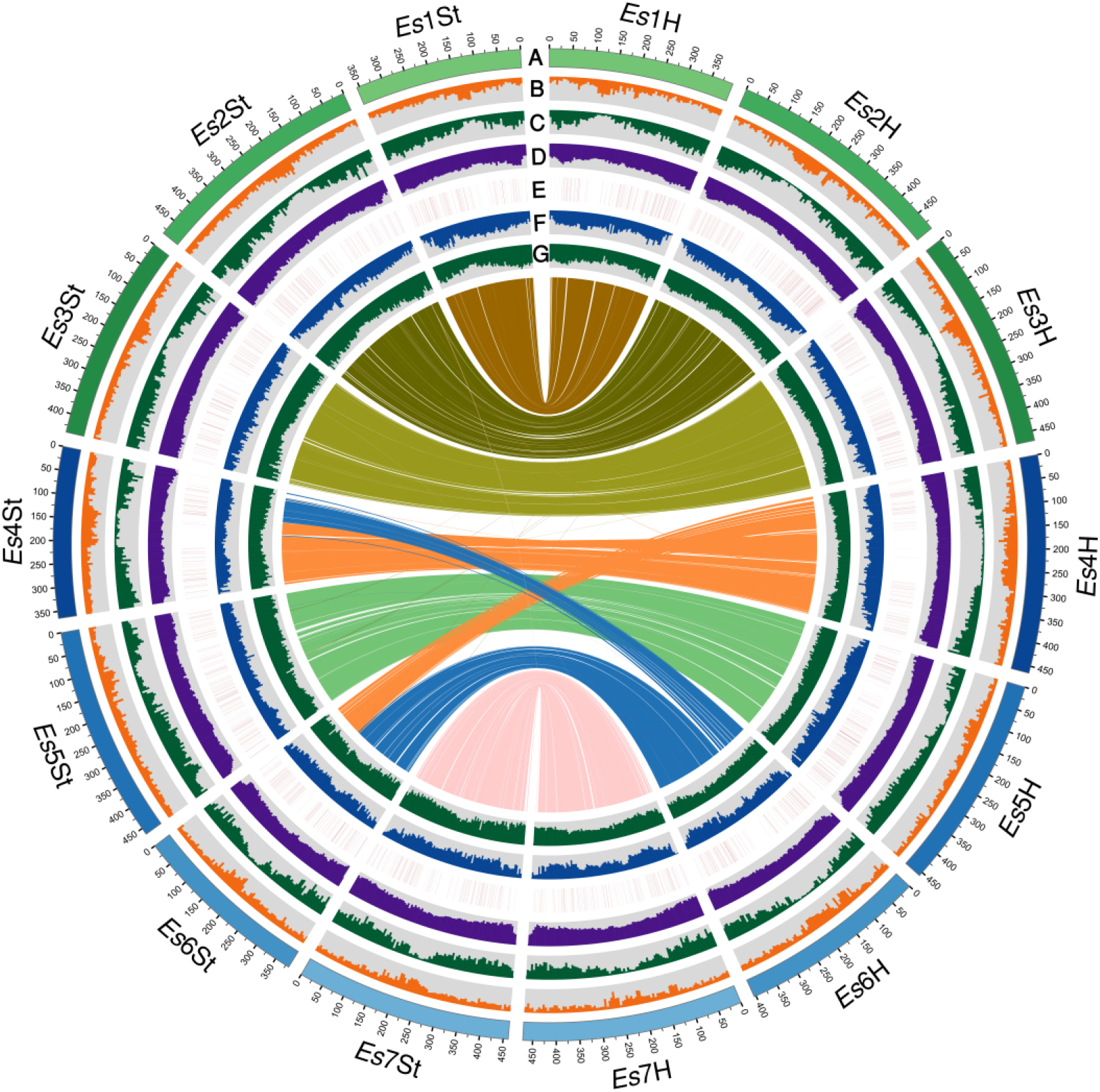
Genomic features of *E*.*s*. Circos display of important features of Genome-wide and synteny between H (*Es*H) and St (*Es*St) subgenomes of *Es* cv. ‘Chuancao No. 2’. The tracks indicate (moving inwards): (A) the 14 pseudochromosomes, (B) the distribution of GC content in each chromosome, (C) the density of gene and (D) transposable elements, (E) the distribution of dominant genes in transcriptome of all samples, (F) the density of SNP and (G) Indel, which was calculated using 5 Mb non-overlap window. The innermost layer shows inter-chromosomal synteny.

**Fig. 2.**
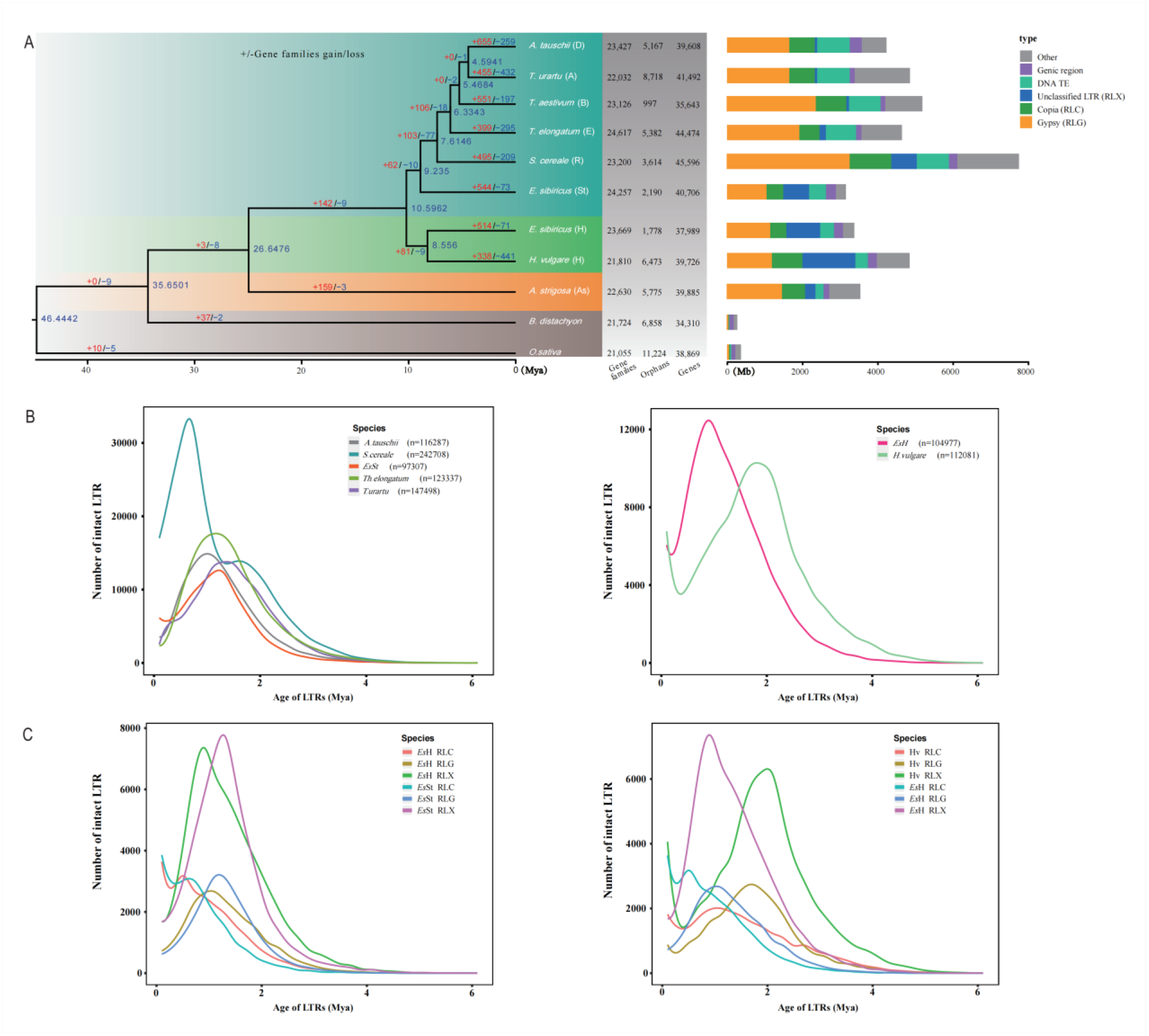
The comparative analysis of genome evolution and genome constituents. (A) A phylogenetic tree and gene family gains (red numbers) and losses (blue numbers) of the *Es* genome and nine grass genomes (left); Genomic constituents in *Es* in comparison with other nine grass species (right). (B) The temporal patterns of LTR-RT insertion bursts in the subgenomes of *Es* and other related species. (C) The insertion bursts of Gypsy, Copia, and unclassified retrotransposon elements between *Es*St and *Es*H, and *Es* and barley. The numbers of intact elements used for this analysis are provided in parentheses.

**Fig. 3.**
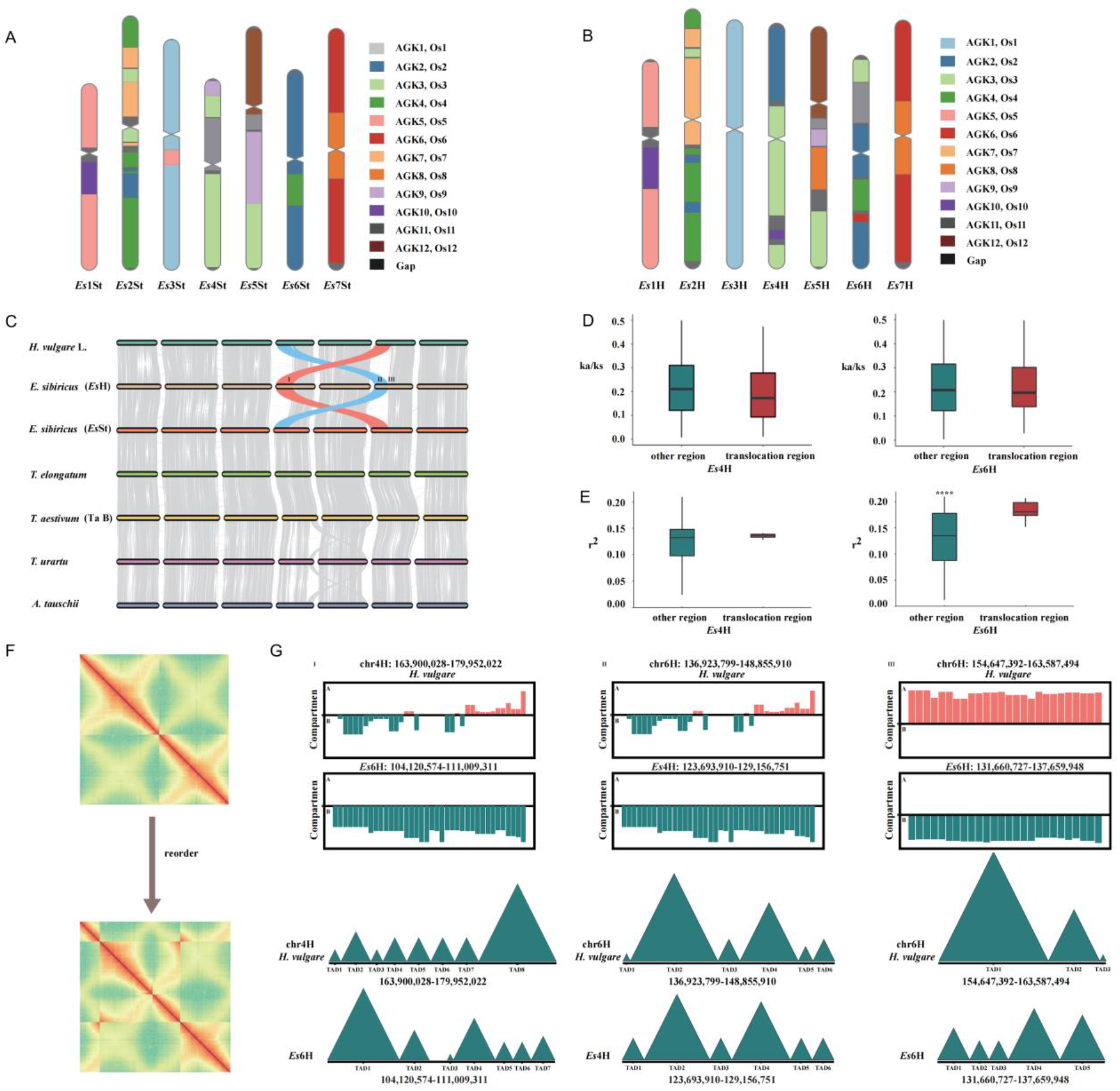
Chromosome evolution, synteny and subgenome dominance analyses of the *Es* genome. The inferred evolutionary trajectories of *Es*St (A) and *Es*H (B) according to the AGK model. (C) Chromosome synteny analysis. The translocation fragment is indicated in blue (*Es*4H) and red (*Es*6H). In addition, the symbols ‘I, II, III’ represent the breakpoint left flanking regions (caused by translocations) in *Es*4H and the breakpoint left and right flanking regions in *Es*6H, respectively. Boxplots of ka/ks values (D) and squared correlation of allele frequencies (r^2^) within 1 Mb (E) of ‘other’ and ‘translocation’ regions. (F) The Hi-C plots for evaluation of the translocation region located in *Es*4H and *Es*6H. The Hi-C interaction heatmap of translocation region was disrupted (down) compared with the original heatmap (up). (G) A/B compartments and topologically associating domains (TADs) of breakpoint flanking regions. Orange represents the A compartments and blue represents the B compartments. One triangle filled with blue color represents one TAD unit.

**Fig. 4.**
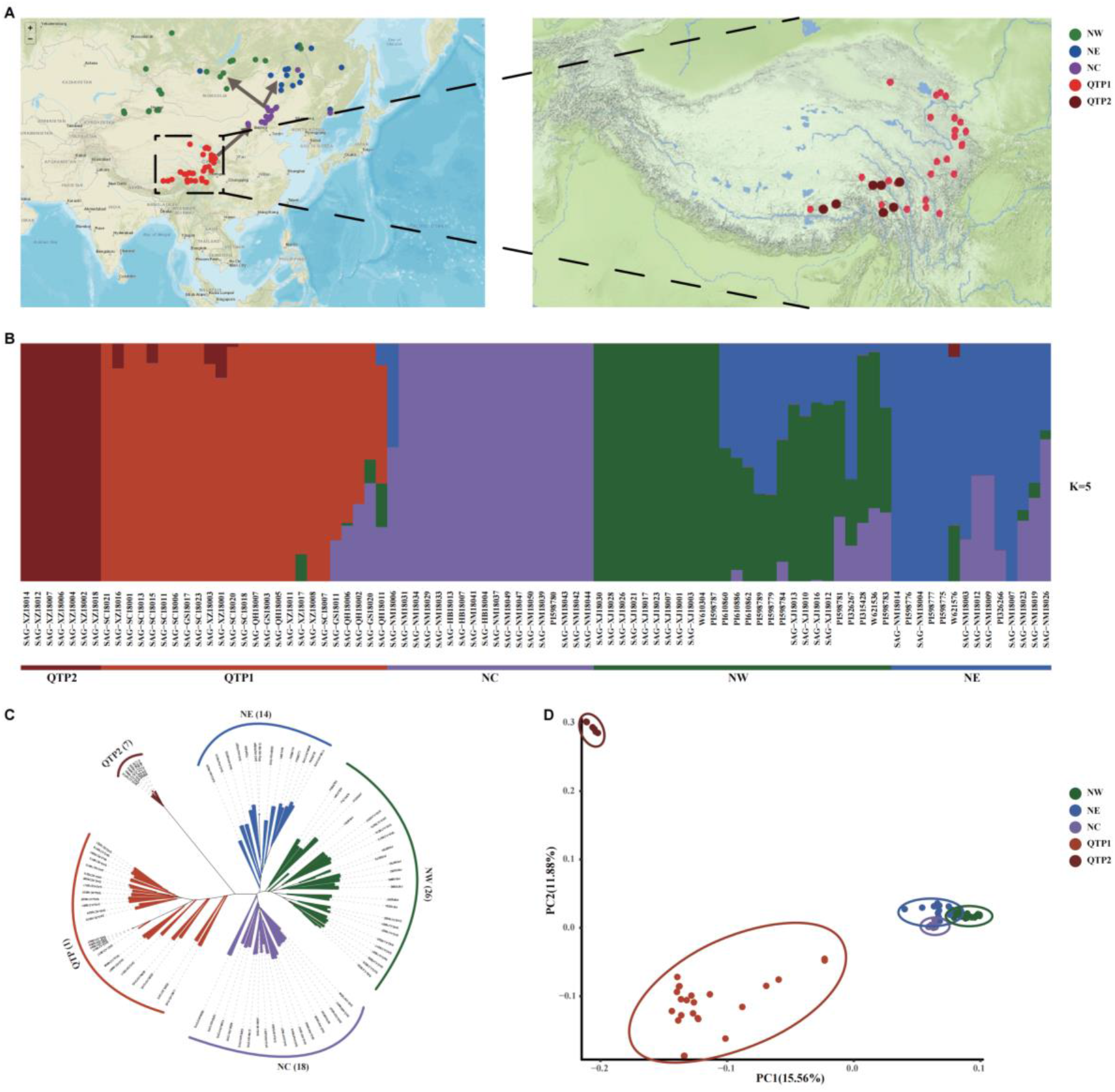
(A) Geographic distribution of the 90 *Es* samples and the detailed geographic distribution of the *Es* samples from Qinghai - Tibet Plateau (QTP1, Light red; QTP2, dark red). (B) ADMIXTURE plot of 90 *Es* samples (k = 5). (C) Neighbor-joining phylogenetic tree of 90 *Es* samples based on SNPs. (D) Principal component analyses plot of the first two components (PC1 and PC2) of the 90 *Es* samples.

### Genome annotation

Our genome annotation identified a total of 101,769 protein-coding genes, consisting of 79,002 high-confidence (HC) and 22,767 low-confidence genes, with around 99.59% of predicted genes having functional annotations (Fig. 1c, Table 1 and Supplementary Table 9, 10). The number of HC genes in *Es*H and *Es*St was 37,989 and 40,706, respectively, and these were considered for further analyses. The average length of genes in *Es* was 6,384.51 base pairs (bp), with 6,270.61 bp in *Es*H and 6,509.49 bp in *Es*St, respectively. Notably, the average gene length of *Es* is significantly longer than in *Triticum urartu* (Tu), a progenitor of the wheat A subgenome ^[18]^ and B subgenome (Ta B) ^[19,20]^, but shorter than in *Aegilops tataschii* Coss (At) ^[21]^, a progenitor of wheat D subgenome. The average intron length of HC genes in *Es*St and *Es*H were 1,242 bp and 1,300 bp, respectively, surpassing the average intron lengths observed in other Poaceae plants (Supplementary Table 11). Additionally, we identified 8,820 non-coding RNAs (ncRNA) in the *Es* genome, including 2,410 transfer RNAs (tRNAs), 1,277 ribosomal RNAs (rRNAs) and 5,133 microRNAs (miRNAs) (Supplementary Table 12).

### Transposable element analysis

In present study, we developed a phylogenetic tree using *Es* (including *Es*H and *Es*St) and nine other sequenced plant species, namely At, Tu, Ta B, *Thinopyrum elongatum* (THe) ^[22]^, *Secale cereale* (Sc) ^[23]^, *Hordeum vulgare* (Hv) ^[24]^, *Avena strigosa* (As) ^[25]^, *Brachypodium distachyon* (Bd) ^[26]^ and *Oryza sativa* (Os) ^[27]^ by 3,025 single-copy orthologous genes (Fig. 2A). The tree revealed that *Es*H and *Es*St formed two distinct homologous clades, with *Es*H closely aligned with Hv, diverging from Hv approximately 8.56 million years ago (Mya). *Es*St clustered with At, Tu, Ta THe, and Sc, with the speciation event estimated to have occurred around 9.24 Mya. Our genome annotation revealed a total of 5.71 Gb transposon elements (TEs), accounting for roughly 86.90% of the *Es* genome. Specifically, 2.95 Gb and 2.72 Gb of TEs contributed to 87.56% and 86.16% to the *Es*H and *Es*St subgenomes, respectively (Fig. 1d, Table 1 and Supplementary Table 13). This was higher than the percentages of At, Tu, TaB, and Hv, which showed 78.24%, 81.16%, 79.83% and 78.16% of the genome sequences, respectively (Fig. 2A, Supplementary Table 14). Analysis of LTR insert time indicated a significant burst of LTRs around 0.9 to 1.2 Mya, a relatively recent event compared with Hv. In *Es*St, a substantial burst of LTRs occurred approximately 1.4 to 1.7 Mya, consistent with the timelines seen in Tu and THe. Despite this, the number of intact LTRs in *Es*St is lower than that in At, Tu and THe, possibly contributing to the smaller size of *Es*St in comparison to these species (Fig. 2B). Furthermore, cross-genome comparisons for genomic composition highlighted that the proportion of Gypsy elements (RLG), Copia elements (RLC) and the unclassified retrotransposon elements (RLX) significantly influenced the genome size (Fig. 2A, Supplementary Table 15). Subsequent analyses of insertion times of RLG, RLC, and RLX in EsH, EsSt, and barley genomes revealed that the insertion times of RLG and RLX coincided with the overall LTR insertion times (Fig. 2C). These results suggest that large-scale bursts of specific retrotransposon superfamilies, particularly RLG and RLX, may directly contribute to the expansion of the *Es* genome, consistent with a previous study on the rye genome ^[23]^.

### Chromosome evolution of *Es*

To elucidate the evolutionary trajectories of *Es* chromosomes, we performed an analyses of the syntenic blocks between *Es*St, *Es*H and the modern rice genome with conservative post-duplication AGK (n=12) intermediate ^[28]^ (Fig. 3A and Fig. 3B). We found that *Es*3H shares a common ancestry with the homologous chromosomes of Tu3, At3 and Hv3, tracing back to a single ancestral chromosome of AGK1 or Os1, indicating a highly conservative evolutionary pattern for this chromosome in wheat tribe ^[29]^. In contrast, its homologous counterpart, *Es*3St, emerged from a nested fusion event involving AGK5 (Os5) and AGK1 (Os1). Both *Es*1St and *Es*1H were formed through a nested fusion of AGK10 (Os10) and AGK5 (Os5), including a translocation of an ancient chromosomal segment from AGK5 (OS5) to the common ancestral chromosome of *Es*1St and *Es*1H. Similarly, *Es*7St and *Es*7H resulted from a common nested fusion of Os6 (AGK6) and Os8 (AGK8), consistent with the observation in Tu7, Ae7 and Hv7 ^[28,29]^. *Es*6St arose from a nested fusion event involving ancestral chromosome Os4 (AGK4) and Os2 (AGK2), distinct from its homologous chromosome of Tu6, At6 and Hv66, which derive from a single chromosome of Os2 ^[29]^ (Fig 3C). Additionally, the *Es*6H homologous to *Es*6St harbored two additional translocations originating from ancient chromosomal segments of Os3 (AGK3) and Os11 (AGK11). Notably, *Es*2St, *Es*2H, *Es*4St, *Es*4H, *Es*5St, and *Es*5H were derived f from complex translocations involving at least three ancestral chromosomes.

The high-quality *Es* genome assembly in this study allowed us to conduct genomic synteny analyses and uncover chromosomal structural variations. As expected, over 90% of genomic regions in both *Es* subgenomes demonstrated strong synteny conservation with other species in the wheat tribe (Supplementary Table 16). We detected 32 translocations and 80 inversions in the *Es* genomes, encompassing 395.93 Mb and 558.60 Mb, respectively (Supplementary Table 17a, b). A notable reciprocal translocation was detected between *Es*4H and *Es*6H in *Es*, spanning approximately 144 Mb on *Es*4H and 112 Mb on *Es*6H (Fig. 3C). This translocation was confirmed by Hi-C links and *in situ* hybridization experiment (Fig. 3F; Supplementary Fig. 5). Moreover, the genotyping analyses for this translocation in 90 *Es* accessions using resequencing data suggests that this inter-chromosomal variation is fixed in the *Es* populations (supplementary Table 18). Importantly, evidence of this translocation was found in 13 accessions of diploid *Hordeum* species, excluding *H. vulgare*, suggesting this variation predates the hybridization of the ancestral StSt and HH donors (supplementary Table 19).

The protein-coding genes within translocation regions displayed lower nonsynonymous/synonymous (Ka/Ks) substitution rates compared to syntenic regions, indicating stronger negative selection on these genes (Fig. 3D). We also analyzed linkage disequilibrium by measuring the squared correlation of allele frequencies (r^2^), within a 1 Mb region flanking the translocation breakpoint on *Es*4H and *Es*6H chromosomes (defined as ‘translocation’) and compared it with other regions (defined as ‘other’). The mean r^2^ values of ‘translocation’ and ‘other’ on *Es*4H chromosome is 0.126 and 0.117 (Wilcoxon rank sum test, *P*=0.641), and 0.195 and 0.146 for ‘translocation’ and ‘other’ on *Es*6H (Wilcoxon rank sum test, *P*=1.539e-05), respectively (Fig. 3E). The higher r^2^ values observed around these breakpoint regions suggest increased recombination rate and reduced genome stability at the exchange loci boundaries in *Es*H subgenome. Furthermore, *Es* genes at the exchange boundaries exhibited apparent A-to-B compartment switching compared to the corresponding regions in barley, indicating alteration in topologically associated domain (TAD) boundaries between *Es*H and barley (Fig. 3G, Supplementary Data 1). Interestingly, genes within the syntenic blocks near the translocation site were enriched in GO terms such as transcription regulator activity (GO:0140110), regulation of cellular macromolecule biosynthetic process (GO:2000112), DNA binding (GO:0003677), and ATP binding (GO:0005524) (Supplementary Data 2). Some of these genes could respond to environmental stresses like cold or drought. For example, *EsiH06g0027800*, encoding a Myb-related protein, responds to both cold and drought stress, and *EsiH06g0027510*, which encodes an Alanine-tRNA ligase, responds to cold stress through the alanine tRNA-mediated pathway ^[30,31]^ (Supplementary Data 3).

### Genetic diversity and population structure analysis

To explore genetic variation in *Es*, re-sequencing was performed on a set of 90 accessions (Fig. 4A and Supplementary Data 4), achieving an average depth of approximately 11.2× and covering about 93.7% of the *Es* genome (Supplementary Data 5). Using this dataset (8.82 Tb), a total of 80,148,422 high-quality SNPs and 11,517,976 InDels were identified (Supplementary Table 20 a, b) through genetic variation calling, amounting to 12.2 SNPs and 1.75 InDels per kb. Of these variants, 1,507,958 SNPs (1.88%) and 150,384 InDels (1.31%) were located within coding regions, with 865,646 SNPs (1.08%) causing codon changes, transcript elongation, or premature stop codons, while 91,093 InDels (0.79%) led to frameshift mutations. Notably, the H subgenome of *Es* exhibited higher nucleotide diversity (π= 7.97×10^−4^) than the St subgenome (π= 7.59×10^−4^). Moreover, the linkage disequilibrium (LD) more rapidly in the H subgenome than in the St subgenome (Supplementary Table 21), suggesting a higher degree of genetic recombination in the H subgenome of *Es*.

The genetic structure of the *Es* populations was assessed for clusters (K) ranging from 3 to 6 based on SNPs data among the 90 *Es* accessions (Supplementary Fig. 6). Both phylogenetic and principal component analyses (PCAs) of the 90 *Es* accessions indicated five clearly major groups (Fig. 4 b, c, d), namely QTP with QTP1 and QTP2 subgroups, NC (Hebei, Inner Mongolia), NE (Inner Mongolia, Russia and Mongolia), NW (Xinjiang, Russia and Mongolia) groups. Notably, seven accessions from QTP formed a unique genetic group (QTP2) in the phylogenetic tree and PCA plot (Fig. 4 c, d), holding a foundational position in phylogenetic trees constructed using potential diploid ancestral species (Supplementary Fig. 7). Conversely, the complex topography and environmental conditions of the QTP have led to numerous glacier refuges ^[32]^, resulting in refuge populations with elevated genetic diversity levels due to their longer history of population dynamics compared to post-glacial populations ^[33,34]^. Nucleotide diversity (π) analyses for the four groups demonstrated that QTP has the highest genetic diversity (π = 7.25 × 10^−4^, supporting QTP as the center of origin for *Es* (Supplementary Fig. 10, Supplementary Table 21).

## Conclusions

A near-complete genome assembly of *Es* cv. ‘Chuancao No.2’ is generated in this study via PacBio and Hi-C data. The 6.57 Gb assembly had an N50 of 4.46 Mb, and more than 99.28% of its sequences were anchored to 14 pseudomolecules. A lot of translocations were found between *Es*St and *Es*H subgenomes, and linkage is closer located in breakpoints and flanking regions of translocation between *Es*4H and *Es*6H. The QTP is one of the centers of origin and genetic diversity of *Es*. These results provide an essential basis for functional genomic research to elucidate the molecular mechanisms underlying important agronomic traits of *Es* and for exploring a wide range of population genetic diversity associated with genome evolution in the future.

## Methods

### Samples collected for genome assembly and resequencing

The *Es* cv. ‘Chuancao No. 2’ from the Sichuan Academy of Grassland Sciences (SAG) was selected for genome sequencing and assembly. Young leaves were collected from an individual plant of *Es* cv. ‘Chuancao No. 2’ in the tillering stage, growing in the germplasm resource park of SAG, located in Hongyuan County, Sichuan Province, China, for DNA extraction, SMRT PacBio, and Hi-C sequencing.

A total of 90 *Es* samples and 25 diploid species from *Hordeum* and *Pseudoroegneria* were collected for resequencing. Among the *Es* accessions, 19 were sourced from the USA National Plant Germplasm System (NPGS), and 71 were extensively sampled from seven provinces in China, covering nearly all major *Es* geographical areas in China. The mature seeds of resequencing samples were collected between July and September 2018 and from 2018 to 2019. Additionally, all 25 diploid species were obtained from the NPGS, with 14 accessions belonging to *Hordeum* and 11 to *Pseudoroegneria*. These seeds were germinated and transplanted to the germplasm resource park of the SAG, Hongyuan County, Sichuan Province, China.

### Sequencing and *de novo* genome assembly

A PacBio SMRT library was prepared and sequenced using the PacBio Sequel II platform. A Hi-C library was created according to the Proximo Hi-C plant protocol, using HindIII as the restriction enzyme for chromatin digestion. Subsequently, the Hi-C libraries were sequenced on the Illumina NovaSeq platform with a read length of 150 bp.

The Canu v1.5 with a correctedErrorRate of 0.025 ^[35]^ was utilized to trim and correct the PacBio subreads. The corrected subreads were assembled using WTDBG (https://github.com/ruanjue/wtdbg). Additionally, the corrected subreads were assembled by MECAT ^[36]^. The MECAT-based assembly served as the reference, and the WTDBG-based assembly was used as the query in a Quickmerge ^[37]^ operation to merge the assembles. Subsequently, the illumina paired-end reads were aligned to the Quickmerge-based assembly using BWA (mapping method: MEM) ^[38]^. Finally, the assembly underwent a polishing step performed by Pilon with parameters --mindepth 10 --changes --threads 4 --fix bases ^[39]^.

The Quickmerged and polished assembly was fragmented into 50 kb fragments, and BWA ^[38]^ was used to align the Hi-C data to the segmented assembly. Only the uniquely mapped reads were preserved. Subsequently, the genome was reconstructed based on these uniquely mapped reads using LACHESIS software ^[40]^, employing the specified parameters: CLUSTER_MIN_RE_SITES = 52, CLUSTER_MAX_LINK_DENSITY = 2, CLUSTER_NONINFORMATIVE_RATIO = 1.3, ORDER_MIN_N_RES_IN_TRUN = 46, ORDER_MIN_N_RES_IN_SHREDS = 42.

### Genome evaluation

The genome completeness was assessed using four methods. Firstly, BUSCO program ^[16]^ was run against the embryophyta dataset with the default parameters, resulting in 97.65% (1576 out of 1614) complete BUSCOs present in the assembly. Secondly, the final assembly was validated by aligning paired-end (PE) short reads with BWA ^[38]^, where 1,316,511,132 PE reads from six libraries were mapped to the assembly at a properly mapping ratio within the range of 98.40% to 98.47%. Thirdly, LAI ^[41]^ was employed to assess assembly continuity using full-length long terminal repeats retrotransposons (LTR-RTs), with LTRharvest ^[42]^ and LTR_FINDER ^[43]^ for *de novo* prediction of candidate LTR-RTs and LTR_retriever ^[15]^ for combining and refining these candidates to obtain the final full-length LTR-RTs. The LAI was calculated based on the formula: LAI = (Intact LTR-RTs length/Total LTR-RTs length) * 100%.

### Repeat and protein-coding gene annotation

A combination method involving *de novo*-based and homology-based prediction was utilized to identify the transposable elements (TE) in the genome of *Es*. Initially, LTR_FINDER ^[43]^ and RepeatScout ^[44]^ were employed for *de novo*-based prediction with default parameters, and the resulting *de novo*-based TE library was subsequently classified using the PASTEClassifier software ^[45]^. Next, the *de novo*-obtained TE library was merged with Repbase ^[46]^ and wheat TE sequences (ClariTeRep: https://github.com/jdaron/CLARI-TE). This combined TE library was then utilized for the annotation of assembly TEs using RepeatMasker (http://repeatmasker.org/). Intact LTR sequences were detected and refined by LTR_FINDER v1.07 (score = 6) ^[43]^. The flanking sequences of the intact LTRs were extracted and aligned using MAFFT with specific parameters (—localpair —maxiterate 1000) ^[47]^. The EMBOSS v6.6.0 software ^[48]^ was employed to calculate the distance “K”. The time calculation was performed using the formula T = K/(2 × r) based on the molecular clock rate 1.3×10^−8^ mutations per site per year.

A combination of ab initio-based, homology-based, and RNAseq-based prediction methods was utilized to annotate the protein-coding genes of *Es*. Ab initio-based predictions employed Genscan ^[49]^, Augustus v2.4 ^[50]^, Glimmer HMM v3.0.4 ^[51]^, GeneID v1.4 ^[52]^, and SNAP ^[53]^. For homology-based prediction, GeMoMa v1.3.1 ^[54]^ was applied using protein sequences from *A. thaliana, H. vulgare, O. sativa, S. bicolor, T. aestivum, Z. mays* as homologous references. RNAseq-based prediction involved transcripts assembly using Hisatv2.0.4 ^[55]^ and Stringtie ^[56]^ with the guidance from the reference assembly. Coding sequences were predicted using GeneMarkS-T (version 5.1) ^[57]^ and TransDecoder v2.0 (http://transdecoder.github.io). Subsequently, the prediction results were integrated using EVM v1.1.1 ^[58]^. Protein sequences were classified as high confidence (HC) if they exhibited an identity of 60% or higher and a coverage of 60% or higher when aligned against protein sequences from *H. vulgare, T. urartu, Ae. Tauschii*, and *T. aestivum*. A total of 79,002 HC genes were identified. These HC genes were then annotated against the SwissProt ^[59]^ and NR ^[60]^ database using BLASTP, and against Pfam (V27.0) ^[61]^ and InterPro (V32.0) ^[62]^ by HMMER ^[63]^ and InterProScan ^[64]^, respectively. KEGG annotation was performed with an e-value cutoff of 1e-5. The Gene Ontology (GO) terms for each gene were derived from the corresponding InterPro or Pfam entries.

### Comparative genomics and chromosome synteny

The protein sequences of *Es, O. sativa, B. distachyon, A. strigosa* (AsAs), *H. vulgare* (HH), *S. cereale* (RR), *T. elongatum* (EE), *T. aestivum* (BB), *T. urartu* (AA), and *A. tauschii (*DD) were aligned using Diamond v0.9.29.130 (http://www.diamondsearch.org/index.php) ^[65]^ with an E-value threshold of 0.001. Subsequently, orthologous and paralogous gene families were identified using OrthoFinder v2.3.7 ^[66]^. These gene families were then annotated using PANTHER V15 ^[67]^. The initial alignment process identified one-to-one single copy gene families. A phylogenetic tree was constructed using IQ-TREE ^[68]^, with the best evolutionary model, JTT+F+I+G4, as identified by ModelFinder ^[69]^ with 1000 bootstrap. MCMCTREE from PAML (v4.9) ^[70]^ was utilized to estimate divergence times with the following parameters: burnin = 5000000, sampfreq = 30, nsample = 10000000. Finally, gene families expansion and contraction analyses were conducted using CAFE v4.2 ^[71]^. To examine chromosome evolution and the structural variations, the study investigated the syntenic relationship between the subgenomes of StSt/HH and *H. vulgare* (HH), *T. elongatum* (EE), *T. aestivum* (BB), *T. urartu* (AA), *A. tauschii* (DD), and *O. sativa*. The analysis was conducted using jcvi software (https://github.com/tanghaibao/jcvi/wiki).

### The 3D chromatin interactions analyses of the translocation site between *Es*4H and *Es*6H

The Hi-C sequencing data from *Es* and publicly available Hi-C data from Hv (ncbi accession numbers: ERR6049631, ERR6049632, ERR6049633, ERR5762911, SRR8922888) were separately aligned to the genome sequences of *Es* and Hv using BWA ^[72]^. Valid interaction pairs and the corresponding standard interaction matrix were verified using HiC-Pro v2.10.0 ^[73]^. Pearson correlation coefficients were calculated, and the resolution of Hi-C data was determined. Normalized interaction matrices at a 100 kb resolution were employed to identify chromatin compartment types by HiTC v1.24.0 ^[74]^, with compartments exhibiting a positive PCA value was designated as A compartments and those with a negative PCA value as B compartments. TadLib (hitad 0.1.1-r1) software ^[75]^ was used to identify TADs at a 20 kb resolution. TADs were considered valid if their Inclusion ratio (IR) value, calculated by HOMER v4.10.1^[76]^, exceeded 0 and their length was greater than 5 bins.

### Variant calling, genetic diversity, and population structure analysis

The clean paired-end reads from 90 resequencing accessions were aligned to the *Es* reference genome using BWA ^[72]^. Picard (http://sourceforge.net/projects/picard/) was employed to eliminate the duplicate reads, and the HaplotypeCaller module of GATK ^[77]^ was utilized to detect single nucleotide polymorphisms (SNPs) and InDels. The preliminary variations were subjected to VariantFiltration in GATK, with the following filtering criteria: QUAL < 30, QD < 2.0, MQ < 40, FS > 60.0, -w 5 -W 10. The high-confidence SNP and InDel variants that passed the filtering were annotated using SnpEff ^[78]^. Finally, the snplist was refined by applying an Integrity threshold of > 0.8 and a minor allele frequency (MAF) threshold of > 0.05, which served as the input for subsequent population genetic analyses.

We utilized Vcftools to estimate genetic diversity indices, including Fst, π, θw, and tajima’D for each group of *Es* accessions. This analysis was conducted in 100-kb windows with a 10 kb step size ^[79]^. Regions with the top 5% of the *F*st and π ratio were identified as candidate selective sweep regions. Principal component analysis (PCA) was performed using EIGENSOFT software ^[80]^. Admixture software ^[81]^ was employed to analyze population structure, and the optimal number of clusters (K) was determined by the lowest error rate during cross-validation. LD decay statistics were computed for various populations using PopLDdecay3.26 (https://github.com/BGI-shenzhen/PopLDdecay) with a parameter setting of MaxDist at 1000 kb.

