## Supplementary figures and images for "A high-continuity and annotated reference genome of allotetraploid Siberian wildrye (*Elymus sibiricus* L., Poaceae: Triticeae)"

### Supplementary Fig. 1.png

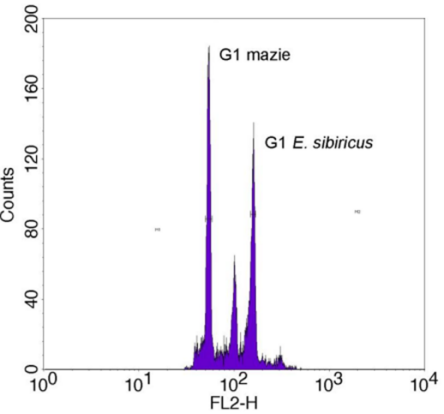

### Supplementary Fig. 2.png

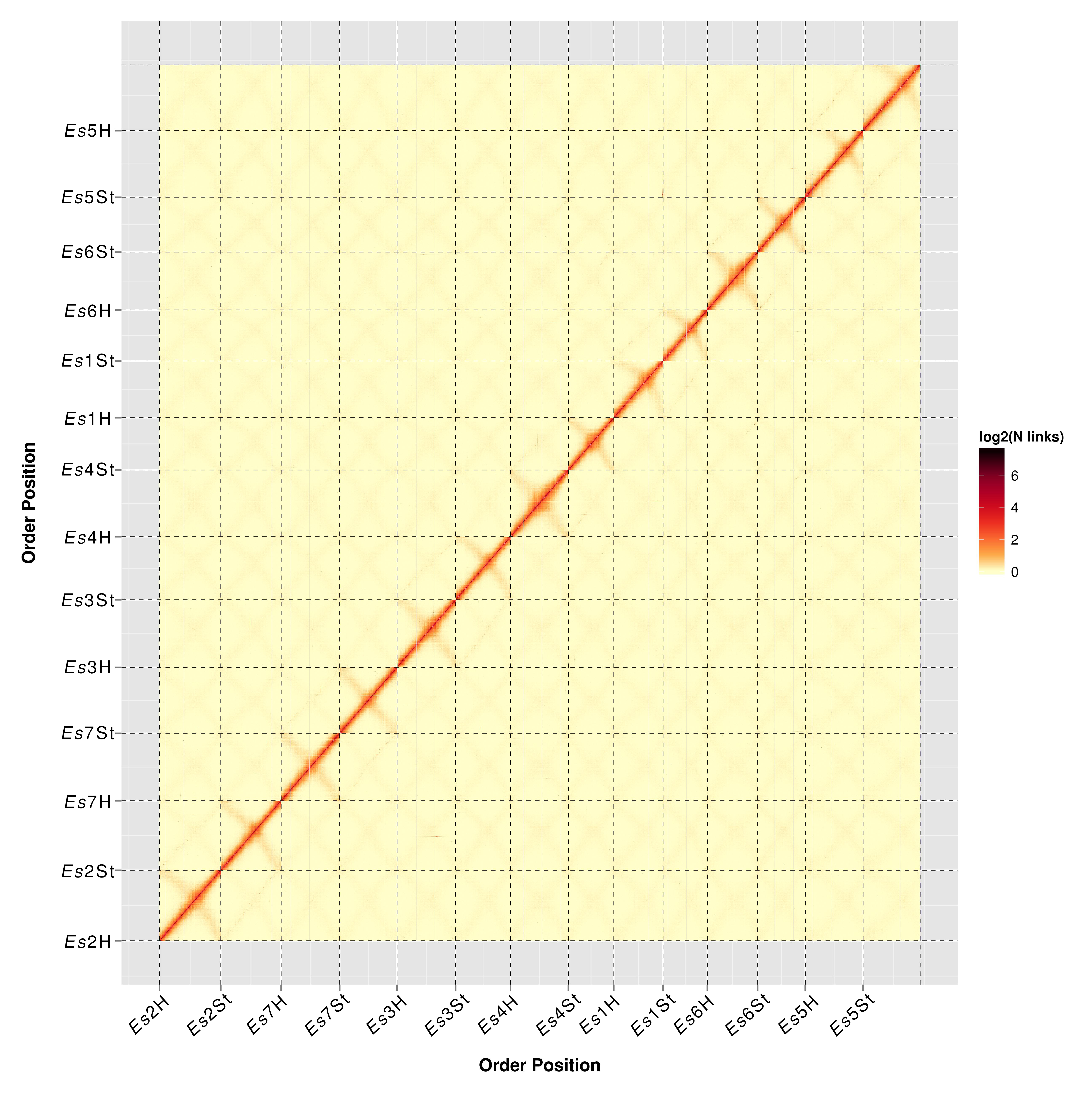

### Supplementary Fig. 3.pdf

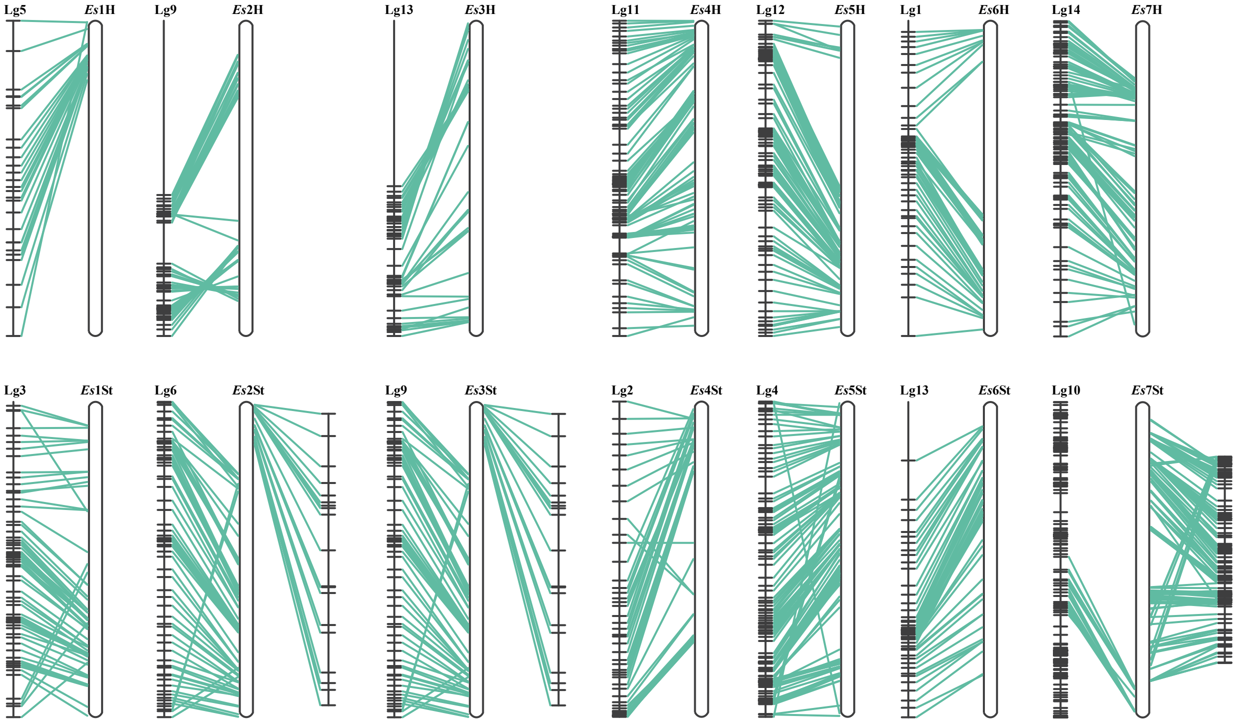

### Supplementary Fig. 4.pdf

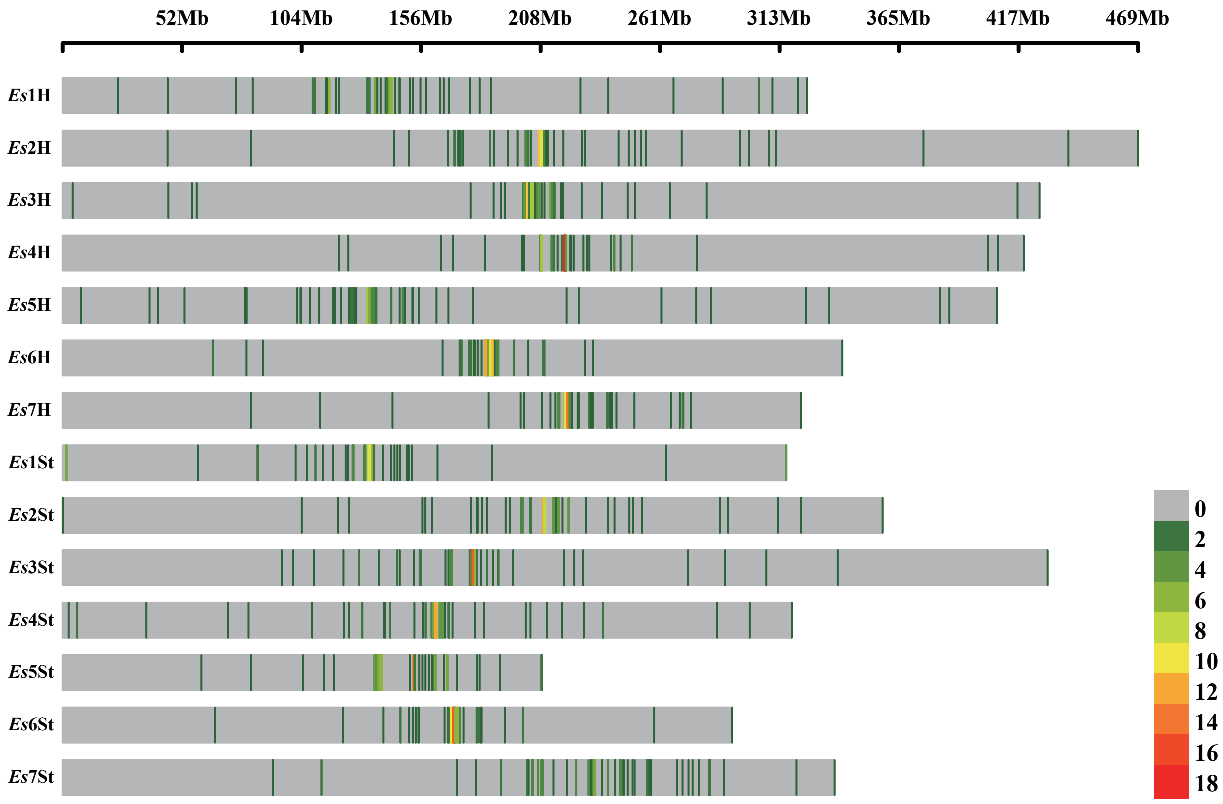

### Supplementary Fig. 5.jpg

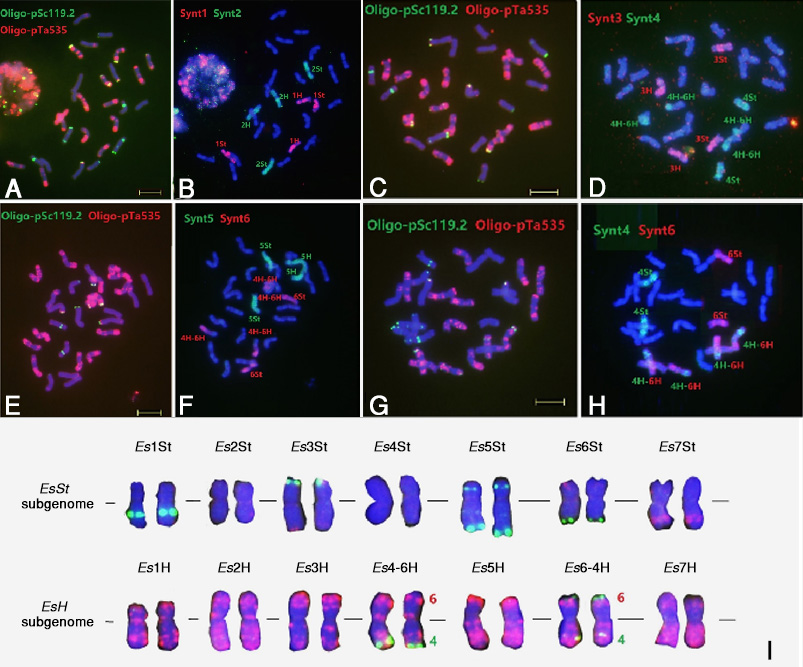

### Supplementary Fig. 6.pdf

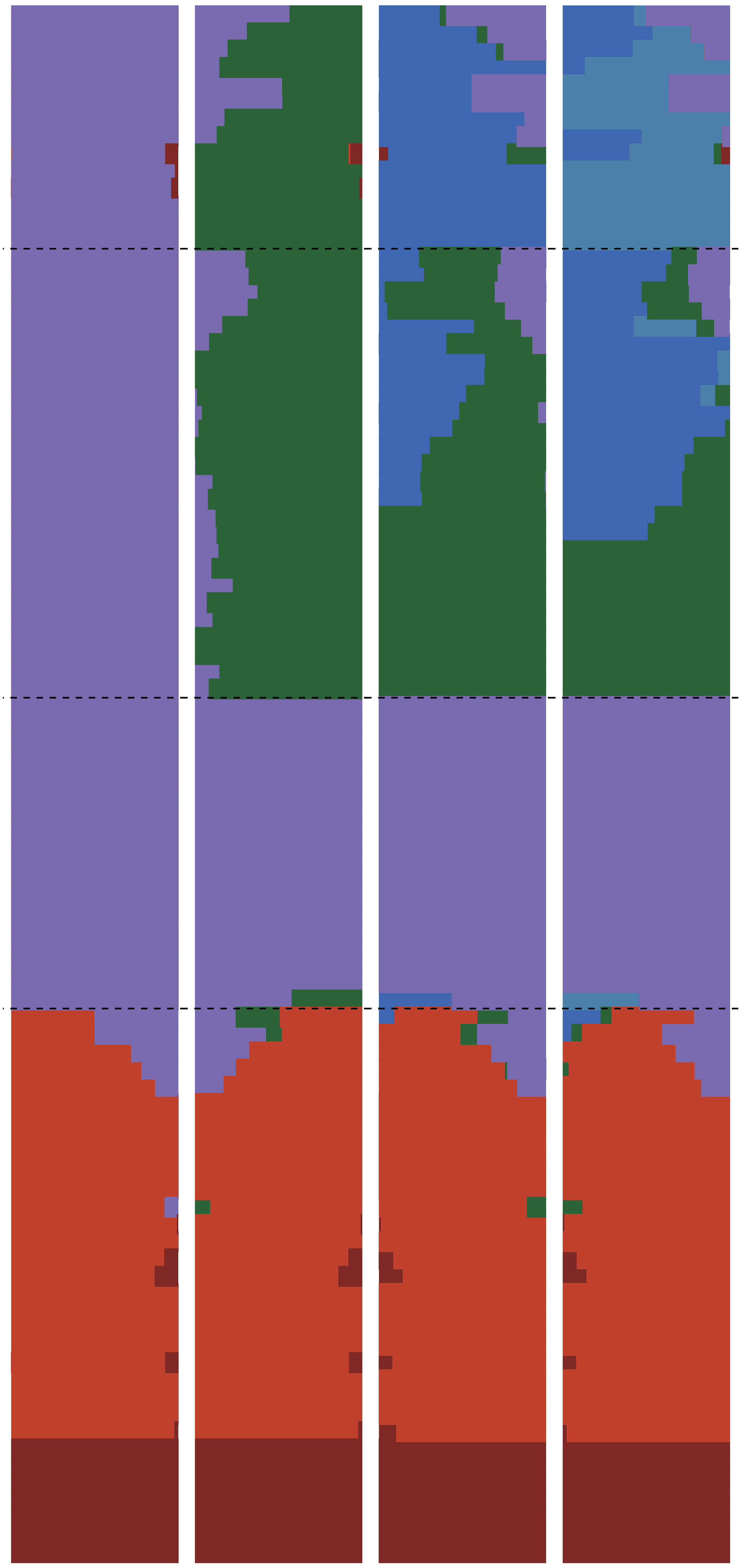

### Supplementary Fig. 7.pdf

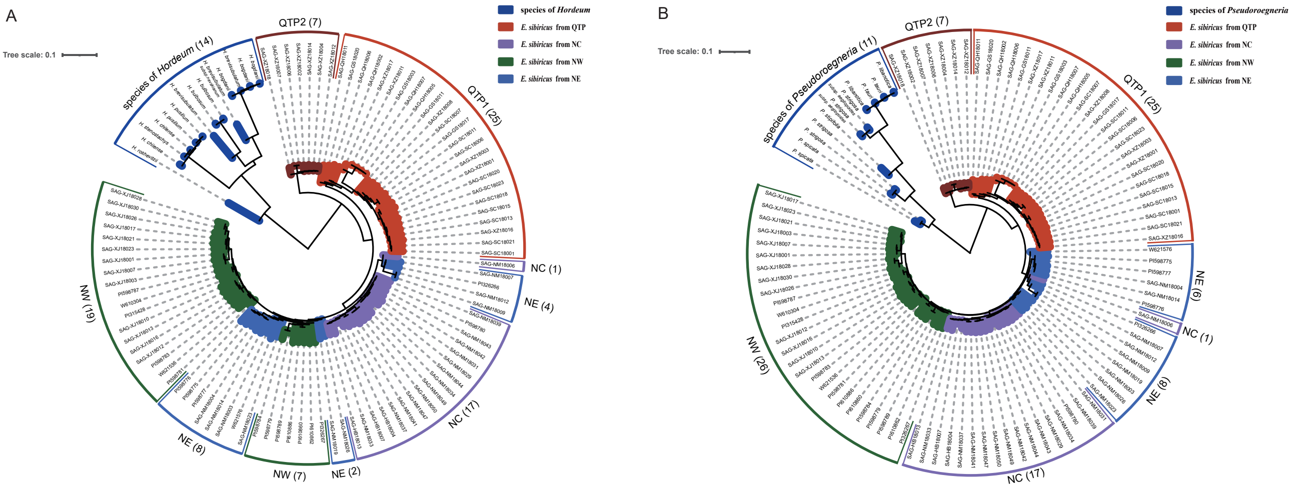
